# Diagnostic assessments of student thinking about stochastic processes

**DOI:** 10.1101/053991

**Authors:** Michael W. Klymkowsky, Katja Koehler, Melanie M. Cooper

## Abstract

A number of research studies indicate that students often have difficulties in understanding the presence and/or the implications of stochastic processes within biological systems. While critical to a wide range of phenomena, the presence and implications of stochastic processes are rarely explicitly considered in the course of formal instruction. To help instructors identify gaps in student understanding, we have designed and tested six open source activities covering a range of scenarios, from death rates to noise in gene expression, that can be employed, alone or in combination, as diagnostics to reveal student thinking as a prelude to the presentation of stochastic processes within a course or a curriculum.

## Introduction

Randomness can be considered from a number of distinct perspectives (Eagle, 2005). A truly random event would be one that has no cause and so would be completely unpredictable. Such events would be expected to display no regularity what so ever, no matter how many similar events were considered, assuming that similar events occur. Such events can be referred to as “acts of god” (miracles) and cannot, by their very nature, be studied scientifically, a point made explicitly in Thomas Paine’s (1794) The Age of Reason. In contrast, there are processes that, while unpredictable at the level of individual events are well behaved (predictable) at the level of larger populations. The presumption is that such processes, commonly referred to as stochastic, are due to as yet unknown, unknowable, or uncalculateable causes. While individual events are independent of one another, individual measurements can be added together in order to arrive at a predictive model for the behavior of a larger population, a behavior known as the “law of large numbers” (Metz, 1998; Tversky and Kahneman, 1971).

There are two generic types of predictably random processes. The first are based on the effects of external factors while the second are due to internal factors. Brownian motion is an example of an externally driven stochastic process. The erratic movements of a molecule or particle at the microscopic level are driven by theoretically deterministic but pragmatically unknowable molecular level collisions.^1^ Our ability to predict the outcome of these molecular level processes is limited by the large number of individual events involved and, in some cases, theoretical limitations such as the Heisenberg Uncertainty principle. While the exact path of a particle exhibiting Brownian motion is unpredictable, the bulk movement (net flux) of large numbers of such particles (diffusion) can be predicted accurately using Fick’s Law (Berg, 1993; Ozawa, 2003).^2^ This seeming contradiction was resolved by Boltzmann and involves the consideration of entropic factors and probability (see Lebowitz, 1993). It was this insight that led Einstein to use an analysis of Brownian motion to establish the existence of atoms (and molecules) (Einstein, 1905; Einstein and Infeld, 1938).

The second general class of stochastic events involves internal factors. This is often presented in terms of a “drunken walk” or hidden variable perspective. The stochastic nature of the process is presumed to be due to factors within the object (the walker). The radioactive decay of unstable isotopes is an example of this type of process (Kossert and Nähle, 2014)^3^, a phenomena apparently not well understood by students (see Prather, 2005).

In the case of such stochastic behaviors a key concept is an event’s probability. We can predict the behavior of populations based on probabilistic models without a detailed understanding of the underlying processes responsible for their behavior. Much of the past research on peoples’ development of an awareness of stochastic processes has focused on when the ability to distinguish deterministic from probabilistic behaviors initially appears (Piaget, 1974; Piaget and Inhelder, 1975). In fact, when, or rather whether, people come to develop an accurate understanding of stochastic processes, so that they can apply such an understanding appropriately to specific scientific scenarios or personal experiences, remains unclear (Metz, 1998).

An obvious question then is, why does an understanding of stochastic processes matter? In biological systems the generation of mutations is a ubiquitous stochastic process involved in evolutionary, developmental, and disease processes (see Baillie et al., 2011; Ionita-Laza et al., 2009). New molecular level techniques have made it possible to move beyond bulk measurements to examine and appreciate the functional significance of the various stochastic behaviors displayed by subcellular systems (genes), individual cells, cellular networks, and social systems (Weber and Buceta, 2013). Such stochastic processes influence gene expression (e.g. Elowitz et al., 2002; Vilar et al., 2003) through their effects on the initiation of transcription and translation, leading to processes such as transcriptional and translational bursting (Eldar and Elowitz, 2010; Jia and Kulkarni, 2011; Pedraza and Paulsson, 2008; Suter et al., 2011), DNA repair (Uphoff et al., 2016), the assembly and disassembly of macromolecules (which together influence molecular half-life), molecular movements, and reaction kinetics. At the cellular and organismic levels, stochastic processes underlie social behaviors such as quorum sensing, altruistic programmed cell death (Engelberg-Kulka et al., 2006; Yarmolinsky, 1995), and the emergence of other phenotypic differences in organisms with identical genotypes, including the appearance of slowly growing drug resistant “persisters”, multicellularity, and the differentiation of stalk and spore cells in *Dictylostelium* (Elowitz et al., 2002; Engelberg-Kulka et al., 2006; Huettenbrenner et al., 2003; Strassmann et al., 2000; Travisano and Velicer, 2004; Uphoff et al., 2016; You et al., 2004).

There are many implications associated with a failure to appreciate the role of stochastic processes in biological systems. Resistance to the role of random mutation as the basis of inheritable phenotypic variation, the ultimate stochastic event in biology, was a major barrier to the acceptance of evolutionary theory; models favoring various deterministic internal or external drivers, including Lemarckian processes, orthogenesis, and/or divine intervention were originally preferred as being more plausible (Bowler, 1992; Bowler, 2005). As an aside, it is in the context of the creation and flow of information that the Central Dogma, as elucidated by Crick (1970) is important; information, generated through mutation and selection, flows out from DNA rather than entering DNA from the environment or some other source (Bowler, 1992; Koonin, 2015).^4^ As noted by others, students often do not explicitly grasp this fact (see Speth et al., 2014 and references therein); similarly students do not appear to grasp the role of genetic drift and associated processes in allele loss, and conventional instruction has been reported to increase their confusion (Andrews et al., 2012). The failure to explicitly consider the origins of biological information is likely to impact student (and teacher) acceptance and understanding of naturalistic (non-theistic) evolutionary processes (Moore, 2008).^5^

There are a few obvious reasons for the difficulty in accepting a role for stochastic processes, most importantly people do not easily accept the role of random or stochastic effects in their own day-to-day lives, rather they prefer active drivers such as fate or luck, or supernatural interventions (Taleb, 2005). A classic example involves what is known as the Gambler’s Fallacy (Turner, 2000; Tversky and Kahneman, 1974), a mistaken assumption that independent events are in fact dependent on one another.

A second issue is that all too often students are not explicitly introduced to stochastic processes during the course of instruction. Andrews et al (2012) note that students are often taught about genetic drift in the context of scenarios (Hardy-Weinberg equilibria) in which drift is theoretically impossible. Similarly, while the specificity and stability of a molecular interaction is based on energy released upon complex formation, the mechanisms behind molecular dissociation are rarely explicitly discussed in the molecular biology textbooks we have examined. Similarly, it is commonplace to find that when molecular level processes are illustrated, particularly in video animations, their stochastic nature is rarely explicitly depicted or acknowledged.^6^ Based on student responses to questions on the Biology Concepts Instrument (BCI)(Klymkowsky et al., 2010) as well as a number of other studies, it is clear that students routinely fail to recognize the role of stochastic processes in biological systems (FIG. 1)(Andrews et al., 2012; Garvin-Doxas and Klymkowsky, 2008; Odom, 1993; Speth et al., 2009; Speth et al., 2014) (Champagne-Queloz, et al, in preparation). For example, the ratio of unproductive to productive collisions within a cell (or in an *in vitro* system) between nucleotide triphosphate molecules and the DNA replication machinery (or the RNA synthesis complex), or between amino acidn charged tRNAs and the mRNA/ribosome complex (during polypeptide synthesis) is generally presented as zero (that is, no unproductive collisions are illustrated), while the actual ratio is typically greater than 4 to 8 for DNA replication (depending on whether one considers deoxy- and ribo-nucleotides) and greater than 20 for polypeptide synthesis (a number complicated by the fact that different amino-acid-tRNAs are present in different concentrations and different codons can be used to direct the addition of the same amino acid).

In the same vein in the context of evolutionary processes, it is uncommon to see any consideration given to the effects of population size, whether from founder effects or population bottlenecks, together with subsequent genetic drift, on the presence of specific traits or a population’s evolutionary trajectory(see Blount et al., 2008; Keinan et al., 2007; Lynch and Conery, 2003). Similarly, it is our impression that few courses consider the stochastic behaviors that influence gene expression and their effects on cellular behavior (see Cottrell et al., 2012; Dar et al., 2015; Eldar and Elowitz, 2010; Jia and Kulkarni, 2011; Raj and van Oudenaarden, 2008; Swain et al., 2002; Uphoff et al., 2016; Vilar et al., 2003). This includes the recent discourse on the role of random events (luck)(Tomasetti and Vogelstein, 2015) versus environmental drivers in the origins of human cancers (Wu et al., 2015).

**Fig. 1:**
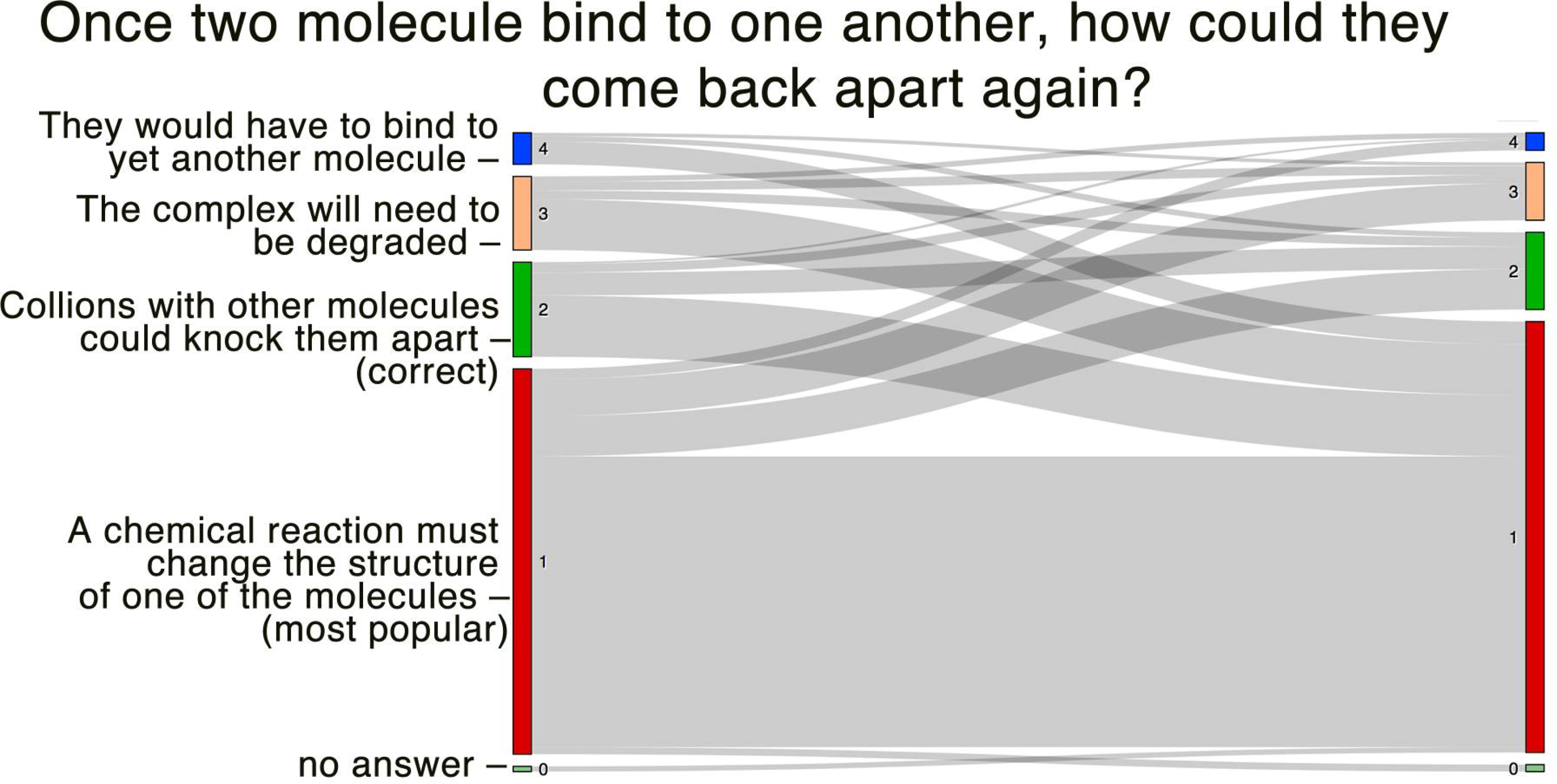
In this cohort of students, there is little change (over the course of instruction) in the recognition that the dissociation of a molecular complex is driven by collisions with surrounding molecules (from Champagne-Queloz, A. Ph.D. et al., in preparation).

As part of a general effort to consider course and curricular content in chemistry and biology (see Cooper and Klymkowsky, 2013a; Cooper and Klymkowsky, 2013b; Klymkowsky and Cooper, 2012; Klymkowsky et al., 2010) and informed by student responses to various BCI questions (Garvin-Doxas and Klymkowsky, 2008; Henson et al., 2012; Klymkowsky, 2007)(unpubl. obs.)(FIG. 1) we have developed tools to gauge students’ understanding of a range of stochastic processes. We use student-drawn graphs and models (Bryfcyzynski et al., 2012; Cooper et al., 2014; Trujillo et al., 2012; Williams et al., 2015) to extend insights gathered through multiple choice and open response questions with the goal of providing instructors and course designers with a clearer picture of student thinking in this area. This effort also reveals aspects of students’ ability to convey their thoughts and construct arguments, as well as their ability to generate and interpret graphs (general numerical and analytic literacy). Two of these activities (death and radioactive decay) involve intrinsic sources of stochastic behavior and four (motion, molecular dissociation, genetic drift, and noise in gene expression) reflect external drivers. Designed as diagnostics, these open source activities can be freely adapted to serve as formative assessments. For each activity, we describe what correct answers should contain and illustrate common students responses we have observed.

**Methods**: These activities were developed and tested using the beSocratic system running out of Michigan State University. To gage student responses, the activities were completed by students in a revised introductory biology course at UC Boulder that emphasizes both evolutionary processes and the qualitative analysis of biological systems; volunteers were awarded 5 extra credit points (added to their final course grade, based on 500 possible points) without regard to whether their responses were correct or not, although completion of all activities was considered in assigning points. Course grades were not curved, so student participation did not negatively impact students who chose not to participate. This activity was judged exempt (IRB protocols 0304.09 and 15–0347). Students’ responses led to revisions of the activities, primarily the deletion of some pages and the consolidation of questions asked.

**Fig. 2:**
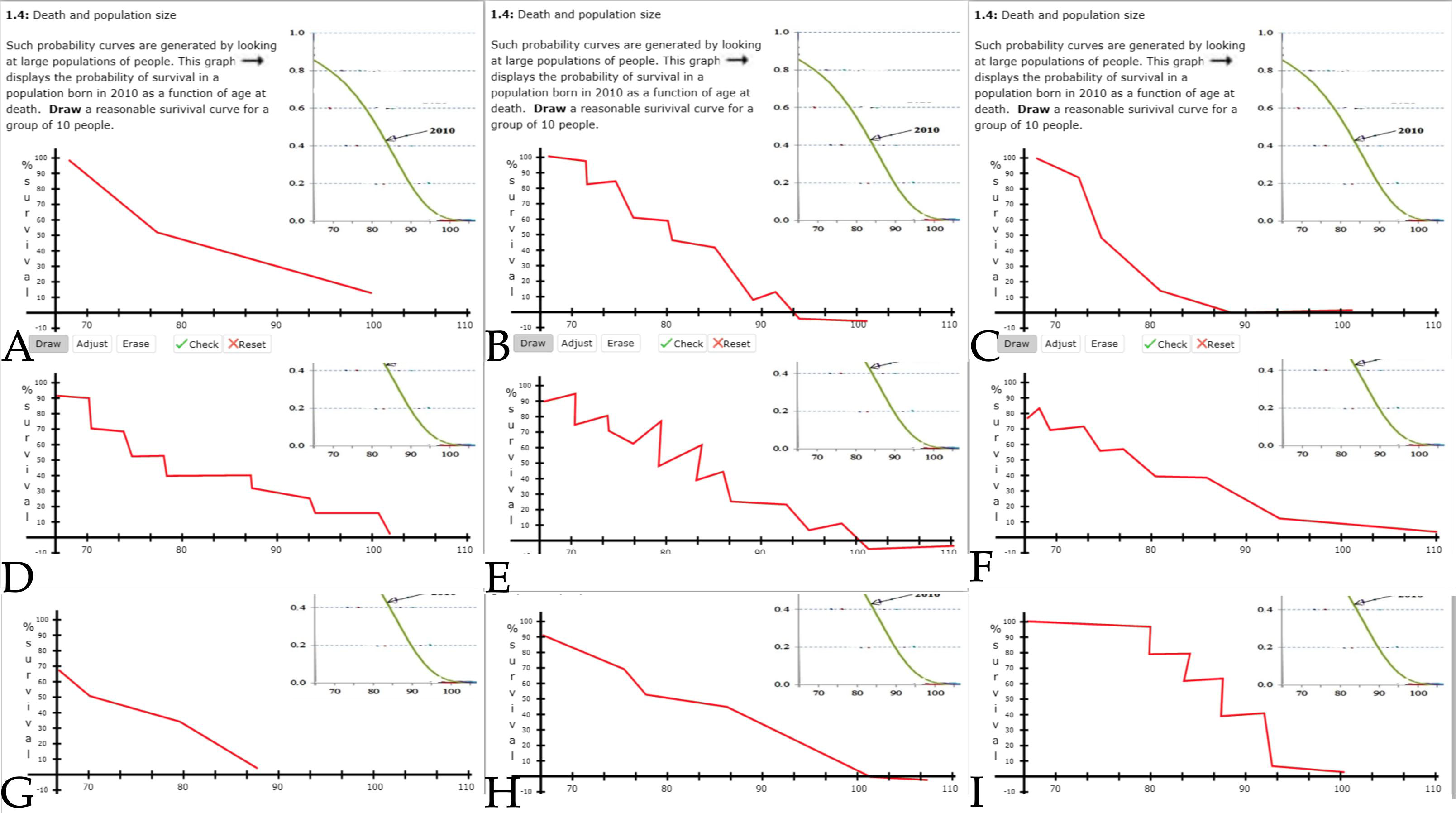
Student drawings of the survival behavior of a small group of 10 people. Often only a few (as illustrated in parts B, D, E, F, and I) recognize that the graph is a step function and that different groups will generate different graphs.

**Description of activities and student responses:** All of the activities begin with a question or two about the student’s educational background, that is, what courses they have taken previously. The activities are typically between 6 to 8 panels in length; students are supplied with necessary background information, multiple choice questions to be answered, and tasks to be completed. Students are generally asked to explain their decisions using open text responses.

**Activity #1 - Predicting death (6 panels):** There are a number of reasons why the ability to predict the probability of death in a population is useful, most notably in the context of insurance. Such institutions accumulate money from their subscribers and, after a certain age, pay out to their members a set stipend. Being able to accurately estimate the average life expectancy of individuals within a population is necessary for the financial viability of such institutions. For a large enough population life expectancy data has the form of a smooth (well-behaved) curve, that is the percentage of the original population that remains alive (or has died) as a function of time is predictable. In contrast, when we consider the fate of smaller groups of people (or of a particular person), the curve has the form of a step function subject to stochastic fluctuations; in the abstract, when a particular person dies is a stochastic event. If students have not previously been introduced to considering the probabilities of events, they will require explicit instruction on probability and its relevance in particular situations. The details of this activity are presented in supplemental data 1.1.

**Outcomes:** Students are expected to be able to recognize, and predict, the differences between the behaviors of large populations (regular and predictable) and small populations (subject to stochastic variation), as witness by their drawing of step graphs which differ between different small groups (FIG. 2). Based on the shapes of survival curves as a function of time, students should be able to deduce that survival rates are influenced by changing environmental factors while the maximum lifespan is not, suggesting that maximum longevity is constrained by inherent (genetic) factors. Students are expected to consider and accurately plot data (something that, perhaps surprisingly, students have great difficulty with), and to recognize how observations of small groups can, by adding them together, be used to make accurate predictions about the behavior of larger groups.

**Fig. 3:**
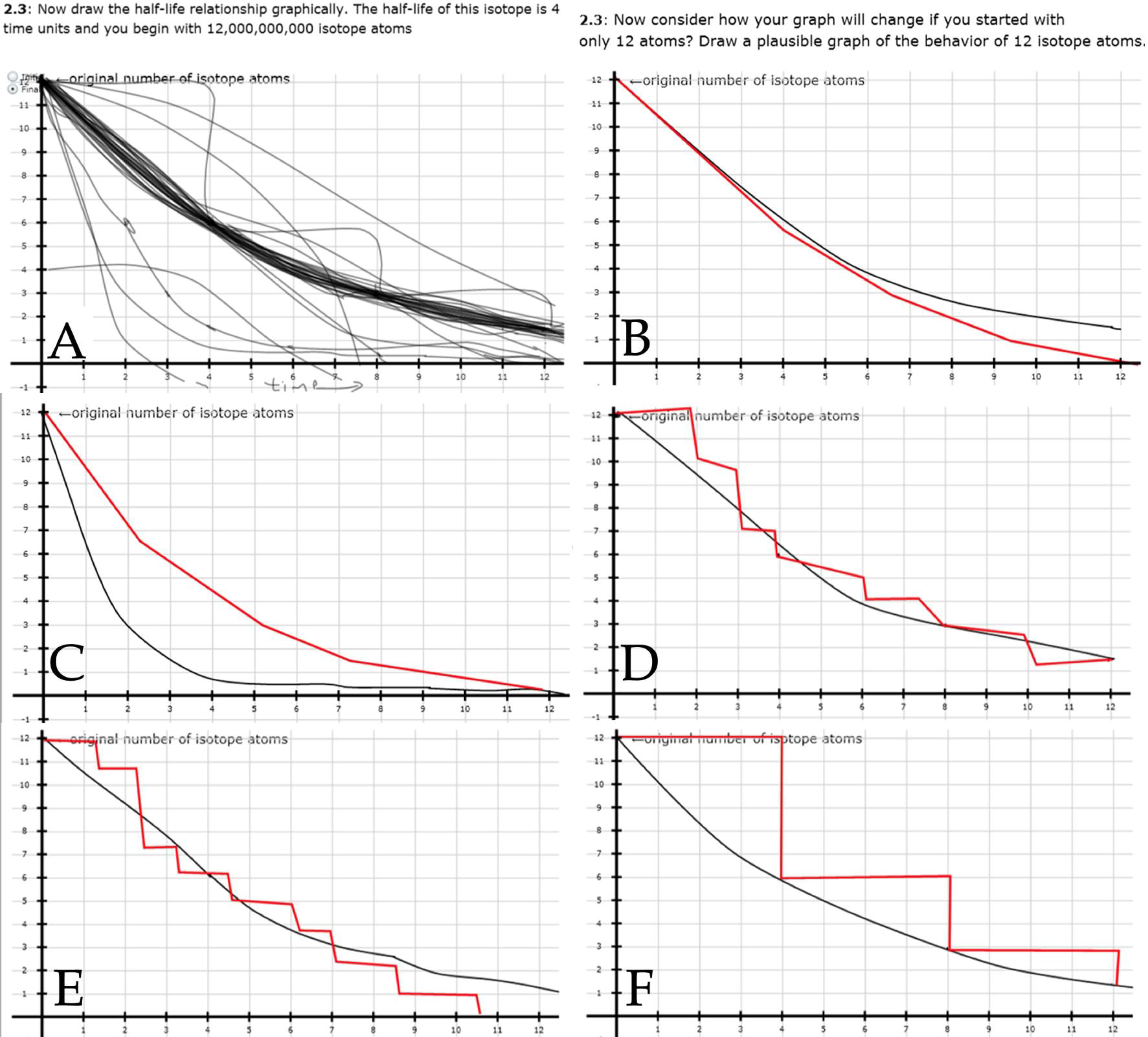
Student drawings of the radioactive decay of a large group of atoms (A: all student decay graphs presented together: parts B-F graphs of individual students - large group decay behavior (in black lines) compared to the behavior of a small group of 20 atoms (red lines in parts B-F). Typically only a few students (illustrated in parts D, E, & F) recognized that the graph of small population behavior is a step function, and that different small groups of isotope atoms will generate different graphs.

**Activity #2 - Predicting radioactive decay (6 panels):** The radioactive decay of isotopes is perhaps the classic example of a stochastic process that can be considered at the level of individual atoms or large populations of atoms. It is likely that students have been introduced to the process of radioactive decay in the course of their education, which provides the instructor (and student) an opportunity to monitor the effectiveness of these educational experiences. If students have not previously encountered isotopes, isotope decay, or the concept of half-life their instructor will need to decide what level of background instruction is appropriate and realistic. We note that the concept of half-life (which applies to a wide-range of biomolecules) is generally not well understood by students (see Prather, 2005). The details of this activity are presented in supplemental data 1.2.

**Outcomes:** Students are expected to be able to predict the differences in the behavior of large and small populations of unstable isotope atoms (FIG. 3). Students are also expected to be able to predict how best to use studies of small populations to predict the behavior of larger groups. This will almost certainly require the presentation of an event’s probability as a function of time.

**Fig. 4:**
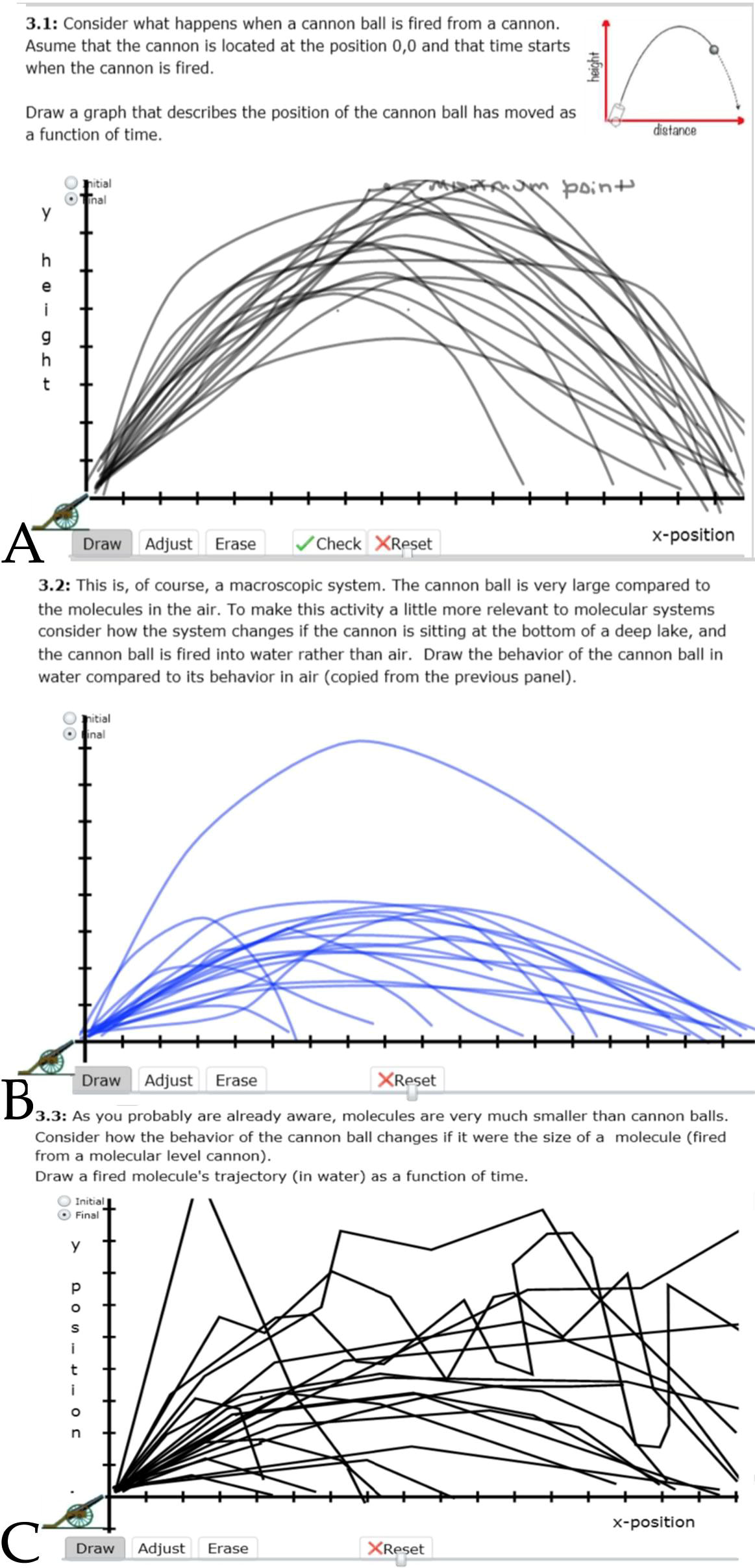
Student drawings of projectile motion in air (A), water (B), and as the size of the projectile approaches the size of a molecule (C). At the molecular level (C) few students recognize the Brownian behavior of molecules.

**Activity #3 - Predicting molecular trajectories (6 panels):** In the context of biological systems, the kinetic movements of molecules supply the energy needed for reactions to proceed (by overcoming the activation energy), for larger molecular complexes to move (diffuse) within the cell, and for complexes, once formed, to come apart again, a critical feature of dynamic systems. However, in most physics courses students are introduced to macroscopic behaviors, such as the movement of objects, based on the concepts of force and momentum. At the molecular level, the movement of particles will be affected by interactions and collisions with other molecules. Indeed, Einstein used a consideration of Brownian (microscopic) motion as evidence for the existence of molecules (Einstein, 1905; Einstein and Infeld, 1938). This formative activity was designed to reveal evidence about whether students can transfer their understanding of motion and interactions from the macroscopic to the molecular level. If students have not been previously introduced to the intermolecular interactions and collisions and the phenomena of Brownian motion, they will likely require instruction on these topics. However, since most students have been exposed to these topics, this activity will provide feedback on whether they are able to apply these ideas. The details of this activity are presented in supplemental data 1.3.

**Outcomes:** Students are expected to be able to generate plausible projectile trajectories in different contexts (e.g. air versus water) and then predict how projectile size (macroscopic versus molecular) influences movement (FIG. 4). This includes recognizing the transition to stochastic (Brownian) behavior as projectile size approaches the molecular. Students are also asked to consider how they might combine observations of small groups to make predictions about the behavior of larger groups.

**Activity #4 - Predicting molecular association and dissociation (8 panels):** Interactions between molecules underlie the binding interactions involved in a range of biological activities. The forces associated with electrostatic attractions are a consequence of the transient (London Dispersion Forces) and permanent dipoles associated with atoms and bonds. The strength of these intermolecular interactions is influenced both by the size and the shapes of the interacting molecules, and the interactions are overcome by the kinetic energy delivered through collisions with other molecules. Responses to this activity provide insights into students’ understanding of the factors that influence molecular association, binding specificity, and dissociation rates. Intermolecular interactions are often considered in the context of introductory (general) chemistry courses, since they explain physical properties such as melting and boiling points and solvent solubility; they are, however, often not well mastered by students (see Williams et al., 2015). If these ideas have not been well understood, their introduction or review may be necessary. The details of this activity are presented in supplemental data 1.4.

**Fig. 5:**
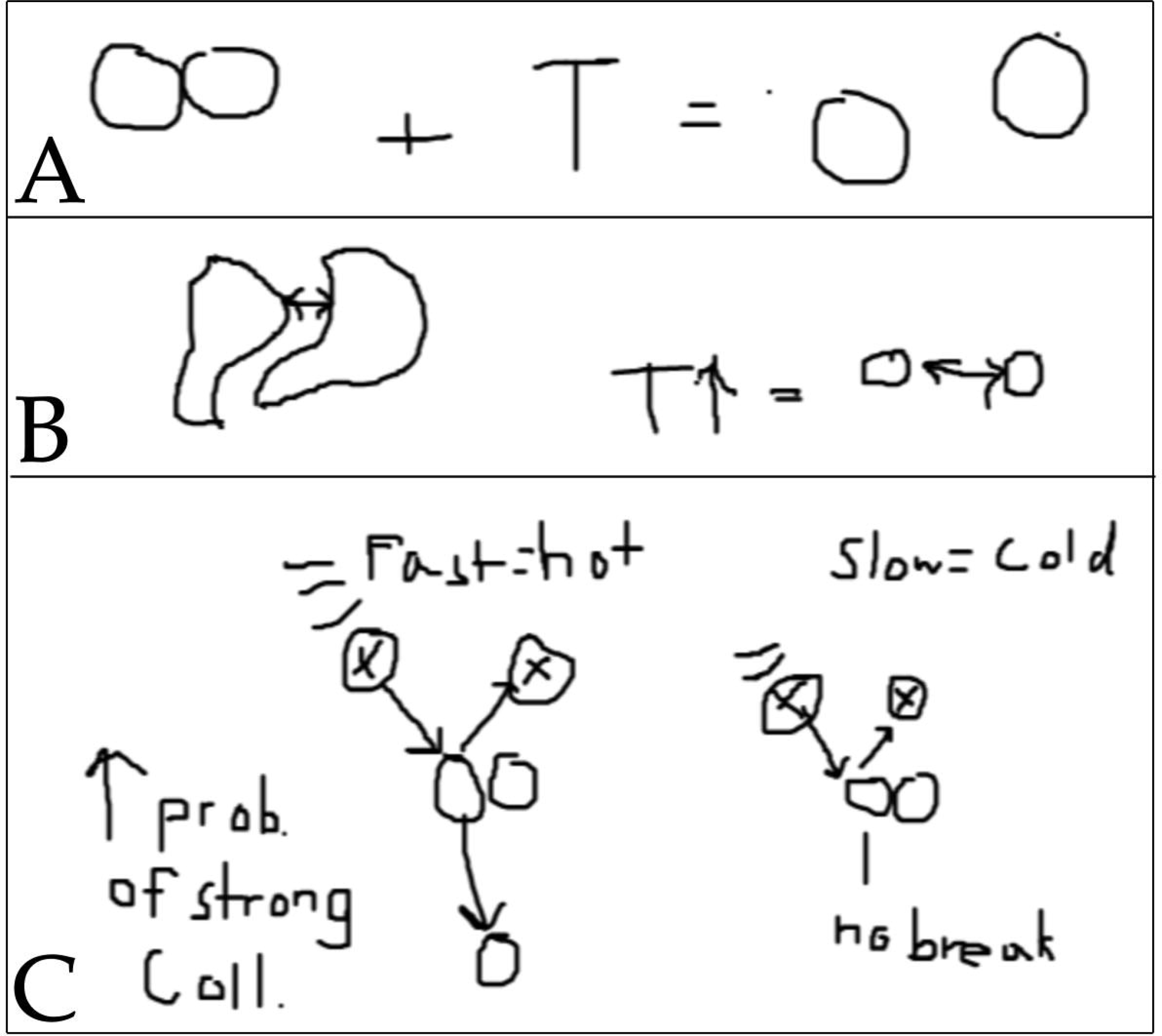
Students’ drawings of molecular dissociation can represent temperature abstractly (A,B) or more directly (C) in the context of molecular collisions.

**Outcomes:** Students are expected to be able to predict (and explain the logic behind their decisions) the relative strength of interactions between molecules, based on their shapes. They are expected to recognize i) how energy is redistributed when molecules interact and to explain their logic in terms of binding (potential) energy, kinetic energy (delivered through collisions with surrounding molecules), and energy conservation; ii) the relationship between kinetic energy, molecular mass and velocity, iii) the distribution of energy in a population of molecules as a function of temperature, and iv) how changes in temperature (typically) influences the stability (dissociation) of a molecular complex (FIG. 5). Entropic effects, associated with the – TΔS term in the free energy equation, are ignored here.

**Activity #5 - Genetic drift (8 panels):** Alleles are generated by the stochastic process of mutation (as opposed to various forms of site directed mutagenesis). Within a population the relative frequency of the alleles at a particular genetic locus is the result of various adaptive (natural, sexual, and social selection) and non-adaptive (genetic drift, founder effects, population bottlenecks, and gene linkage) effects. In the case of genetic drift, the allele frequencies within a population are influenced by stochastic processes. In sexually reproducing species, these processes include which alleles are delivered to which gametes and which gametes combine to form the next generation.

**Fig. 6:**
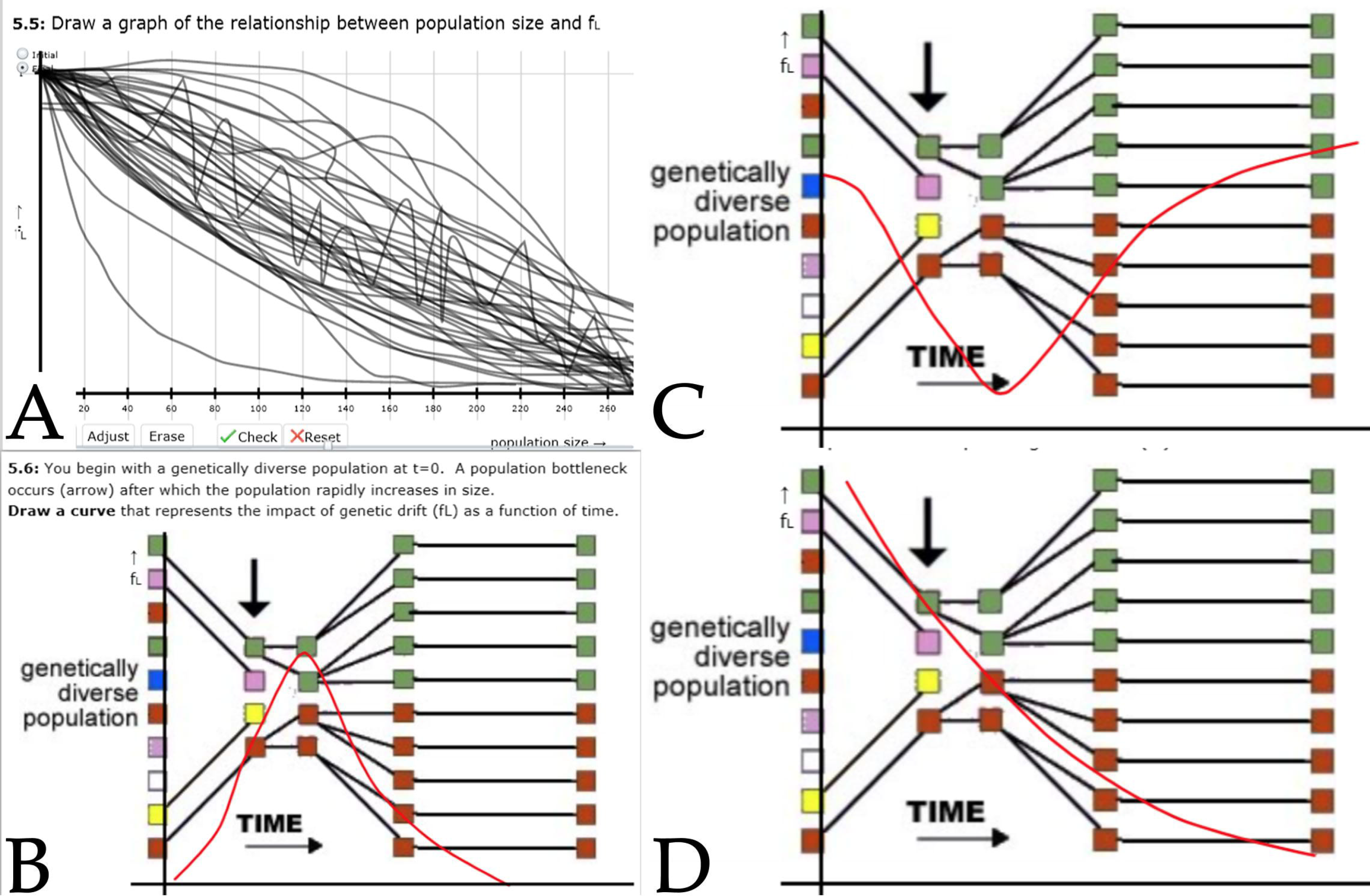
**A**: The ensemble of student graphs (black lines) of F_L_ as a function of population size. Individual student graphs (red lines) of how F_L_changes as a function of a change (a dramatic decrease) in population size. **B** reflects an accurate graph, C indicates a change in F_L_in the wrong direction, while D suggests an alternative (ambiguous) conception.

Our consideration of genetic drift is based on a Java-based web application that can be used by students to generate their own data sets; it does not explicitly consider the processes involved in producing genetic drift.^7^ Because running Java can pose security complications on some computers, we supply students with a data set (obtained from this applet) in the context of the activity. The details of this activity are presented in supplemental data 1.5.

**Outcomes:** The genetic drift activity expects students to be able to describe their understanding of genetic drift and to recognize when an allele as been lost or fixed within a population. Students are expected to be able to calculate the observed frequency of allele loss (F_L_) as a function of population size, to plot that data, and to explain why it makes sense to draw a “best fit” curve rather than connecting the empirically derived values of F_L_. They are expected to be able to predict (based on the relationship between F_L_ and population size) the effects of drastic changes in population size on the value of F_L_ as well as to speculate on the impact of genetic drift on future evolution and to explain their reasoning. In this context, it is worth noting that Andrews et al (2012) reported that distinct misconceptions related to genetic drift not only persisted but increased in frequency after instruction on genetic drift. This provides a rationale for examining student thinking using this or another activity (see Price et al., 2014) in a pre-/post-instruction assessment model.

**Fig. 7:**
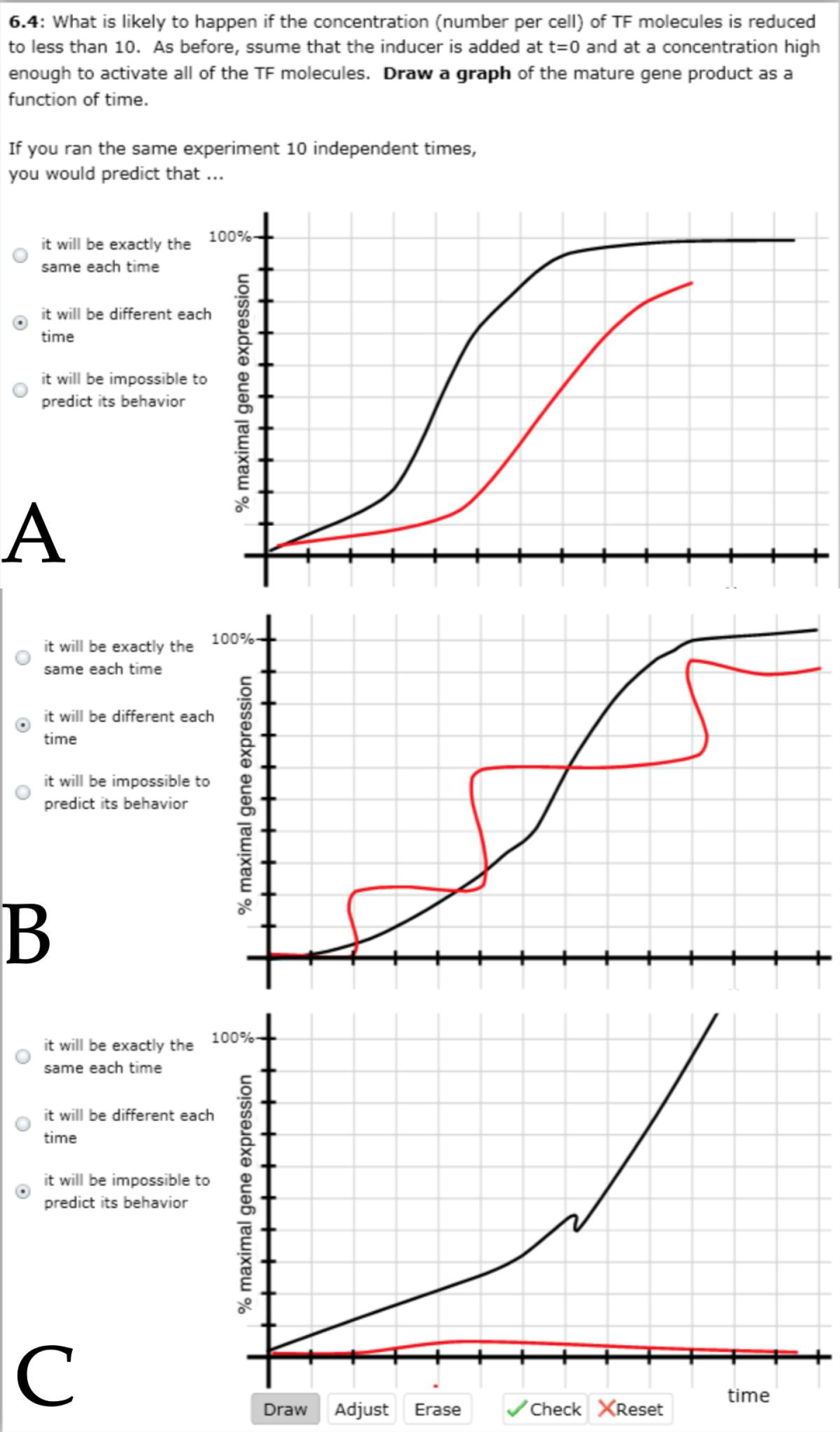
Students are asked to compare (graphically) the effects of high (black) and low (red) concentrations of a transcription factor. **A** depicts similar patterns for both, B depicts a more stochastic pattern of expression, while C suggests that there is no transcription at the low transcription factor concentration.

**Activity #6 -Noisy Gene Expression (5 Panels):** This activity considers the various stochastic factors that influence gene expression. Noisy gene expression, a process that includes transcription, translation, assembly, and localization, is a phenomenon that has at its root in the fact that most cells have only two copies of any particular gene and that the number of molecules regulating the expression of a particular gene, while varying widely, remains relatively small, ranging from less than 10 per cell (in the case of the lac repressor) to many thousands.^8^ Outcomes from this activity can reveal how students approach questions related to individual gene and cell behavior. It is assumed that students already have an understanding of the basics of gene expression, including the processes that influence binding and dissociation (Activity 4) of transcription factors with DNA, as well as the interactions between the mRNA polymerase and DNA (responsible for the termination of transcription), mRNA and splicing factors and the nuclear pore/cytoplasmic transport complex, and the ribosomal complex, including the role of stop codons and release factors in the termination of translation. The details of this activity are presented in supplemental data 1.6.

**Outcomes:** Students are expected to be able to describe (and diagram) the processes associated with gene expression (the binding of transcription factors, the recruitment and activation of RNA polymerase, the opening of the DNA double helix, etc.). They are expected to be able to generate a plausible graphical representation of the appearance of a gene product in a cell that contains a large (10,000) number of the positively acting transcription factors that regulate the target gene, and then a comparable representation of how reducing the number of those transcription factors to 10 will influence their graph. Based on their responses, students rarely explicitly indicate the times required for transcription and mRNA processing to occur. While translation can begin before transcription is complete in prokaryotes, eukaryotic genes often have many large introns that must be removed (spliced out) before the mRNA is transported into the cytoplasm and can interact with the translational machinery. Finally students are asked to predict and justify how the numbers of transcription factor molecules will influence transcriptional noise.

Here are two examples of correct responses to the effect of changing transcription factor concentrations on transcription.

*“The more transcription factors are present, the more likely they will bind to the DNA and begin the process. If there are not as many, the probability of binding will be much lower and the process will be noisier.”*

“*With higher concentrations of TF, the probability of it colliding with the correct sequence increases and therefore decreases the random noise*”.

**Examples of incorrect responses**: *“depends on rna polymerase because that is what creates the gene’s expression. Transcription factors are only in transcription and the beginning of translation”*

“*Higher concentration means that more molecules are colliding and moving around the cell, making more noise.”*

## Discussion

The six diagnostic activities described here can serve as a means to monitor students’ thinking about a range of stochastic processes. It is not our intention that any particular course would use all six activities, since different courses focus on different content areas. That said, all of these processes are deeply analogous, with similar conceptual foundations; they offer an opportunity to determine whether students’ understanding of stochastic processes in one context is transferred to other contexts during a course or within a curriculum.

Based on students’ responses to one or more of these activities, changes in instructional design, both in terms of the content presented and stressed and skills practiced can be considered and their effects monitored. In our own case, it is clear that while the course within which these activities were initially tested sought to improve student understanding of the presence and implications of stochastic processes, there are areas in which further instructional redesign, particularly student time on task and constructive feedback, are needed.

While designed as diagnostics, the activities presented here can be used as teaching tools. For example, while stochastic processes are the common theme, different activities include other important concepts, such as the difference between environmental factors and evolved traits (activity #1), radioactive decay (activity #2), the factors that control molecular association and dissociation (activity #3), molecular movement (activity #4), non-adaptive evolutionary processes (activity #5), and gene regulatory processes (activity #6). A course designer or instructor could adapt these activities to fit the specific context of their course or curriculum. What is critical is that they can reveal lacunae in students’ understanding or ability to apply stochastic processes. While the analysis of textual responses and drawings is, by its very nature, more time consuming, the results of the associated multiple choice questions and student generated graphs can be quickly evaluated and even used as exemplars within a class, while the more time-consuming analyses of students’ textual responses can be used as part of a redesign process, refocusing course goals, materials, and activities. A tangential benefit associated with having students generate and interpret graphs, as well as open text responses is that it can reveal the presence of previously unexpected issues with students’ ability to analyze data, generate and interpret graphs, or to construct logical, empirically-based arguments and conceptual justifications.

Ultimately, the use of these and similar activities can be expected to reinforce the importance of stochastic processes in biological systems, and to help build courses that acknowledge their importance and the conceptual difficulties associated with grasping them in an accurate and useable manner. Ignoring these aspects of biological systems, which has been all too common, encourages a rigid deterministic view of living systems that is directly contradicted by how these systems actually behave and evolve.

## Footnotes

1 https://en.wikipedia.org/wiki/Brownian_motion

2 https://en.wikipedia.org/wiki/Fick%27s_laws_of_diffusion

3 http://phys.org/news/2014-10-textbook-knowledge-reconfirmed-radioactive-substances.html

4 While not surprising given Griffiths’ discovery of bacterial transformation, the discovery of horizontal gene transfer complicates this view.

5 http://www.gallup.com/poll/170822/believe-creationist-view-human-origins.aspx

6 See HHMI: http://www.hhmi.org/biointeractive/translation-basic-detail, http://www.hhmi.org/biointeractive/dna-transcriptionadvanced-detail, DNA Learning Center: https://youtu.be/TfYf_rPWUdY, Harvard Biovisions: https://youtu.be/wJyUtbn0O5Y

7 See http://darwin.eeb.uconn.edu/simulations/jdk1.0/drift.html

8 See the B10NUMB3R5 site: http://bionumbers.hms.harvard.edu

## Acknowledgements

*This work was supported in part through NSF grants DUE-1043707 and 1341987 (M.M. Cooper, PI, MW. Klymkowsky, Co-PI). We acknowledge various ideas supplied through conversations with Natalie Jones (UC Boulder) who helped build a genetic drift activity for the Biofundamentals course.*

